# Discovery of circulating cell-free DNA 5hmC biomarkers for peritoneal metastasis in colorectal and appendiceal cancer

**DOI:** 10.1101/2025.09.18.676945

**Authors:** Yuri F. M. Malina, Lu Gao, Ankit Dhiman, Diana C. West-Szymanski, Yaniv Berger, Xiao-Long Cui, Zhou Zhang, Marco Rivas, Urszula Dougherty, Chang Ye, Akushika Kwesi, Zifeng Deng, Biren Reddy, Hunter D.D. Witmer, Phillip J. Hsu, Chuan He, Wei Zhang, Marc Bissonnette, Kiran Turaga

## Abstract

**Introduction:** Peritoneal metastases (PM) are associated with poor prognosis in patients with colorectal cancer (CRC) or appendiceal adenocarcinoma (AA), yet detection of PM is unreliable using current circulating DNA technology. Leveraging novel 5hmC-seal technology to detect ultra-low amounts of DNA in plasma, we demonstrate the feasibility of 5-hydroxymethylcytosine (5hmC) signatures derived from circulating cell-free DNA (cfDNA) as biomarkers for PM.

**Methods:** Using a highly sensitive and robust 5hmC sequencing approach on genomic DNA isolated from peripheral blood samples, we developed predictive models to identify biomarkers for peritoneal metastases.

**Results:** We obtained genome-wide 5hmC profiles from 71 CRC/AA patients with PM, 41 without PM, and 73 non-cancer controls. Predictive models trained on genomic region 5hmC levels in patients with cancer could distinguish PM status with high sensitivity and moderate specificity (AUC 0.827, sensitivity 92.4%, specificity 46.1%). Pathway enrichment analysis identified epigenetically dysregulated cancer, cell migration, adhesion, and immune-related pathways in PM.

**Conclusion:** Novel 5hmC-Seal technology based 5hmC signatures can detect patients with peritoneal metastases from CRC and AA, albeit with reduced specificity. This study lays a foundation for future clinical assay development for PM.

**Statement of significance:** We demonstrate high-sensitivity detection of peritoneal metastasis in colorectal and appendiceal adenocarcinomas using 5hmC-Seal of plasma cfDNA. Earlier detection of this condition could expand curative treatments in ∼20,000 affected U.S. patients.

## Introduction

Peritoneal metastases occur in 8-13% of CRC cases,^1–3^ and 23% of AA cases,^4^ with an annual burden of ∼20,000 patients in the United States^5,6^. Detection of peritoneal metastases is challenging with low sensitivity of conventional imaging. A meta-analysis found pooled sensitivity of 68% for the detection of PM using CT,^7^ and even lower sensitivity (11%) for early lesions (<0.5 cm).^8^ Surgical exploration by laparoscopy has a higher sensitivity (98%),^9^ but is known to underestimate peritoneal cancer index (PCI) in most cases,^10^ xiphopubic laparotomy is therefore often used as a diagnostic gold standard. The need for surgical procedures for accurate diagnosis leads to inadequate early detection of these patients and consequently inappropriate treatment or exclusion from clinical trials.

Liquid biopsy-based detection tools have shown promise and may offer significant advantages over existing modalities in the detection and management of patients with colorectal and appendiceal cancer.^11,12^ However, both cfDNA and ctDNA technologies have yielded very unreliable results in the detection of peritoneal metastases with sensitivity of <60%, especially in mucinous and appendiceal tumors.^13,14^ This is due to the low quantities of DNA shed by peritoneal metastases as compared to other visceral metastases.

Therefore, there is a critical need for the deployment of robust, ultra-sensitive technology that could facilitate the early detection and treatment of patients with PM from CRC and AA. We developed 5hmC-Seal technology-based on selective chemical labeling and enrichment that requires nanograms of DNA.^15^ This method has been validated to distinguish patients with cancer from healthy subjects in a variety of cancers,^16–21^ and to identify genome-wide 5hmC distributions associated with specific tissues and cell types.^22,23^

Therefore, we hypothesized that PM cells might have a distinct 5hmC signature, and that this signature could be detectable in plasma-derived cfDNA, permitting us to distinguish CRC and AA patients with peritoneal metastasis from those without.

## Materials and methods

### Study population and sample collection

Patients with confirmed CRC, AA and non-cancer (NC) control subjects were consented and enrolled in the IRB approved protocol (IRB10-209A). Primary cancer was confirmed by tissue biopsy in all patients with CRC and AA. Patients with appendiceal mucinous neoplasms were excluded.^24^ Surgical exploration was considered the gold standard to confirm the presence/absence of PM. Matched (age, race, gender) control subjects were recruited from individuals undergoing routine screening colonoscopies without aggressive pathology (adenomas/cancer). Whole blood samples (5 mL) were collected in EDTA Vacutainers from patients prior to colonoscopy (for NC), surgery or laparoscopy (for PM^+^ and PM^-^). Within 6 hours of sample collection, 1-2 mL plasma was isolated from the blood and archived at −80 °C.

### DNA Extraction, 5hmC library construction and high***-***throughput sequencing

DNA extraction was performed using the QIAamp Circulating Nucleic Acid Kit (Qiagen, Cat. No. 55114).5hmC libraries were constructed for all samples using high-efficiency 5hmC-Seal technology.^21^ cfDNA extracted from plasma was end-repaired, 3′-adenylated using the KAPA Hyper Prep Kit (KAPA Biosystems) and ligated with Illumina-compatible adapters. Paired-end 50-bp high-throughput sequencing was performed using a NovaSeq 6000 (RRID:SCR_016387).

### Data processing and quality control

All sequencing data were trimmed using Trim Galore (version 0.6.7, RRID:SCR_011847),^25^ and subjected to quality control analysis using FastQC (version 0.11.9, RRID:SCR_014583).^26^ The results for each sample were collated and analyzed using MultiQC (version 1.21, RRID:SCR_014982).^27^ Samples with at least one of the two paired-end files with fewer than 10M unique reads were excluded from all subsequent analyses due to library preparation and/or sequencing failure. This resulted in the exclusion of 7/192 (3.6%) samples (Supplementary Figure S 1). The reads were mapped to the human genome assembly (GRCh37 hg19) using Bowtie 2 (version 2.4.4, RRID:SCR_016368).^28^

### 5hmC consensus peak set construction and annotation

5hmC peaks were called for each sample “.bam” file using MACS3^29^ (version 3.0.1, RRID:SCR_013291). All peaks were then combined into a merged peak set using DiffBind (version 3.12.0, RRID:SCR_012918),^30^ retaining only peaks occurring in at least 10% of the samples. Raw 5hmC-associated read counts for each sample in the merged peak-set regions were determined using featureCounts (version 2.0.3, RRID:SCR_012919). The resulting merged peak set was further filtered into a consensus peak set by excluding peaks occurring on the two sex chromosomes, peaks less than 5,000 bp in length, and retaining peaks that had ≥15 counts in ≥90% of the samples.

### Region annotation, differential 5hmC enrichment and functional analysis

All regions were annotated using annotatr (version 1.28.0).^31^ 5hmC metagene plot was constructed using deepTools (version 3.5.5, RRID:SCR_016366).^32^ Differentially enriched 5hmC regions (DhMRs) between conditions were determined using DESeq2 (version 1.42.1, RRID:SCR_015687). A combined p-value and log fold change cutoff or an adjusted p-value cutoff was then applied depending on the use case (values specified in figures.) All heatmaps were generated using ComplexHeatmap (version 2.18.0, RRID:SCR_017270),^33^ with all rows and columns clustered using a Ward.D2 Euclidean distance measure.

The 2,000 DhMRs with the greatest absolute value Wald statistic from DESeq2 were analyzed, using rGREAT,^34^ to identify enriched gene ontology terms in the msigdb:C2:CP:KEGG database with a minimum geneset size of 15, using the default algorithm for region-gene assignments. Pathways with a hypergeometric adjusted p-value <0.05 were plotted in a network diagram along with their associated DhMRs using ggraph (version 2.2.1, RRID:SCR_021239).

### Predictive model training and evaluation

All 71,597 consensus 5hmC peaks were considered as candidate features. An elastic net regularization model was then trained and evaluated using nested CV with 10 inner folds and 10 outer folds stratifying samples to maintain similar proportions of sample PM status and sample primary site (e.g. CRC or AA) across folds, repeated 100 times. Class weighting was applied considering only the PM status. Model training was performed using nestedcv (version 0.7.8),^35^ with ranger (version 0.16.0, RRID:SCR_022521) Random Forest (RF) filter.^36^ The top 50 (sparse) or 5,000 (high-accuracy) features resulting from the RF filter were used for elastic net regularization model training.

## Results

### DNA Extraction and 5hmC-Seal assay

After excluding 7 samples for sequencing failure, we included 185 patients, including NC (n = 73) controls and CRC (n = 63) or AA (n = 49) patients (Figure 1). Of the 112 cancer patients, 71 were PM^+^ (63.4%), whereas 41 were PM^−^ (36.6%). Furthermore, 22 (53.7%) PM^−^ patients and 16 (22.5%) PM^+^ patients had primary tumors present at the time of plasma sampling (Table 1).

**Figure 1:**
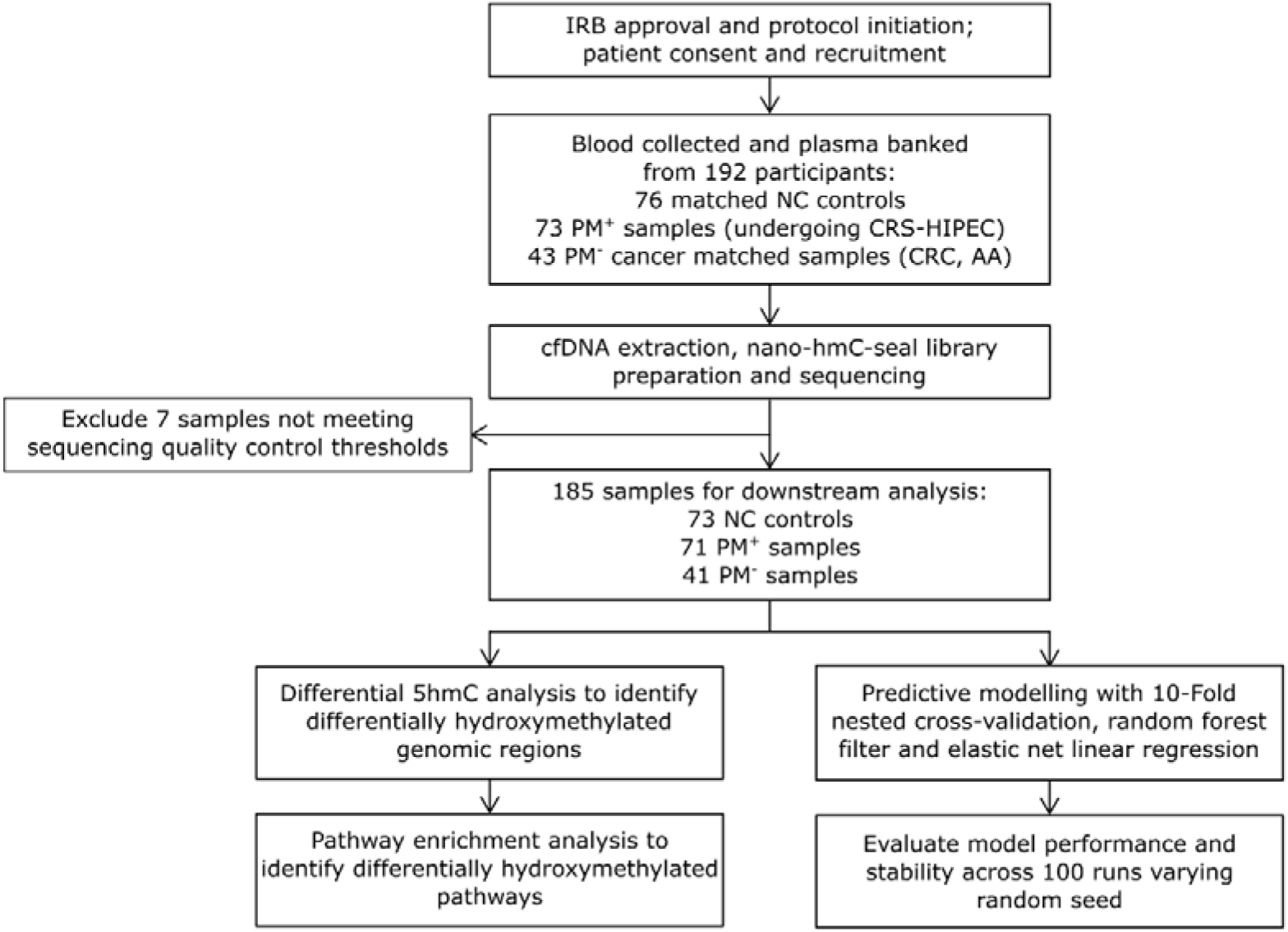
Study design. Graphical outline of the study design. Patients were recruited, and consent was obtained. Blood samples were obtained from 192 participants, and plasma was banked. Cell-free DNA (cfDNA) was extracted, and 5hmC-Seal libraries were prepared and sequenced. Seven samples were excluded for not meeting sequencing quality control criteria. 185 samples (73 NC controls, 71 PM^+^ and 41 PM^-^) were retained for downstream analysis. Differential 5hmC and pathway analysis was performed. Separately, predictive modelling was carried out using a 10-fold nested cross validation approach, combined with random forest filtering, and elastic net regularization regression. NC: non-cancer; PM: peritoneal metastasis; IRB: institutional review board; CRS/HIPEC: cytoreductive surgery with hyperthermic intraperitoneal chemotherapy; CRC: colorectal adenocarcinoma; AA: appendiceal adenocarcinoma; cfDNA: cell-free DNA.

**Table 1:**
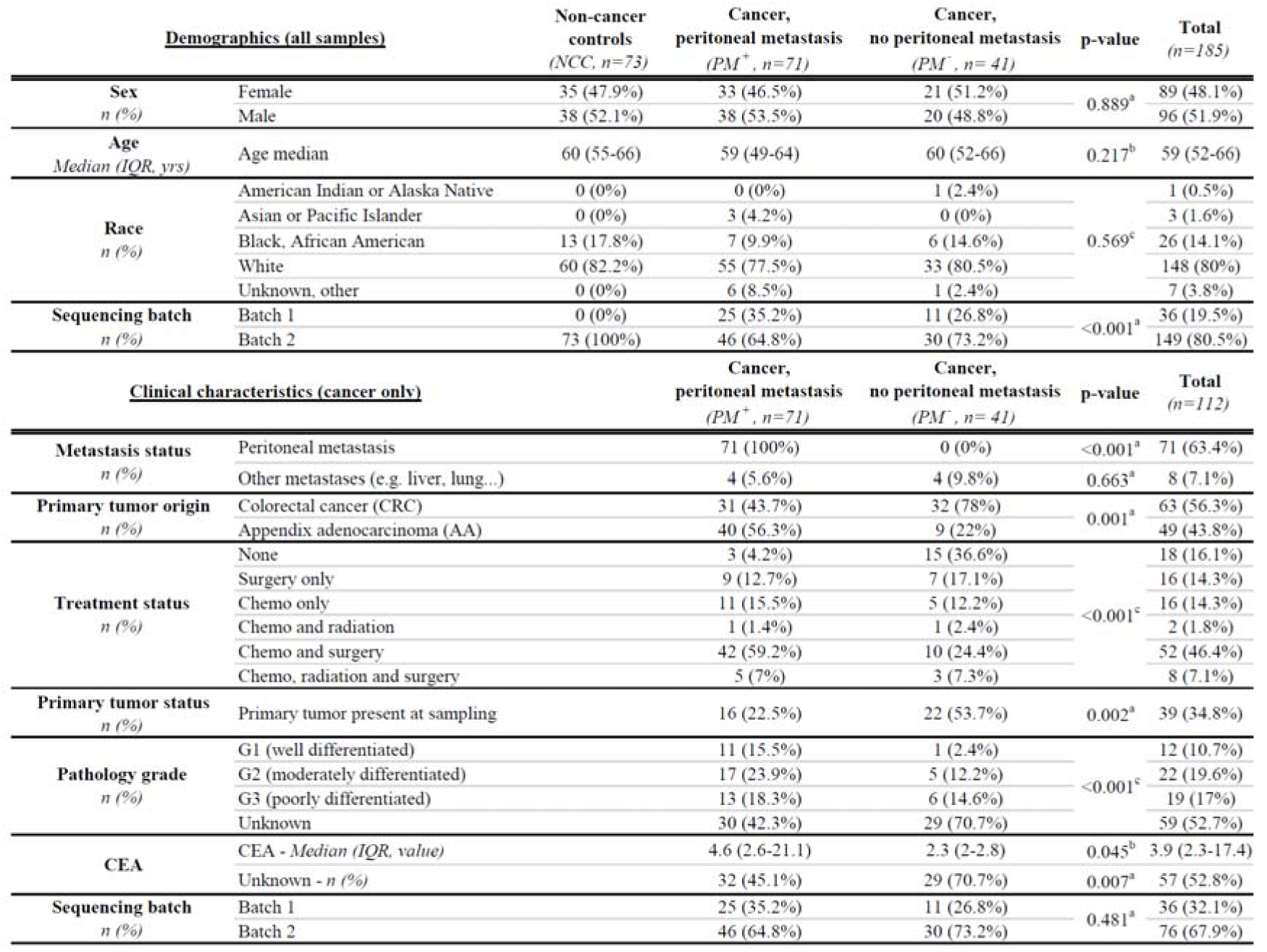
Patient demographics and clinical characteristics. Analysis of participant demographics and clinical characteristics shows that Age and sex are well balanced across conditions. Ethnicity has similar proportion of white and non-white samples, however, is imbalanced when considering other ethnicities (p-value = 0.011). Sequencing batch is imbalanced due to a mix of non-cancer controls and cancer samples having been processed in Batch 2, and only cancer samples having been processed in Batch1. The presence of other metastases and sequencing batch are well balanced across PM conditions. Treatment status and resection of primary tumor prior to sampling are unbalanced as expected due to patients with and without peritoneal metastases following divergent treatment courses. Pathology grade and CEA values are likewise unbalanced as expected due to divergent pathophysiology of metastatic cancer. (a) Fisher’s exact test. (b) Wilcoxon Rank Sum test. (c) Chi-squared test. CRC: colorectal adenocarcinoma; AA: appendiceal adenocarcinoma; CEA: Carcinoembryonic Antigen; IQR: interquartile range. G: grade.

9.7 ng of cfDNA (IQR: 9.7 – 10.0 ng) was extracted from 1 – 2 mL of plasma (Supplementary Figure S 1 a, Supplementary Table 1). Downstream analysis was performed on 63 CRC (34.1%), 49 AA (26.5%) patients, and 73 NC controls (39.5%) (Figure 1). Batch effects identified by principal component analysis were adjusted using ComBat-seq^37^ (Supplementary Figure S1c-h).

### 5hmC profiles in patients with PM^+^

Consensus 5hmC-enriched regions were primarily located within gene bodies (74.4%), and mapped to known enhancer regulatory regions (85.4%, Supplementary Figure S 2 c-d). PM^+^ samples had decreased density of 5hmC peaks in the 5’ UTR than NC controls (6.8% fewer consensus 5hmC-enriched regions per million reads, p-value = 0.037) (Supplementary Figure S 2 a, Supplementary Table 2). Global 5hmC-hypomethylation in cancer samples compared to NC controls is consistent with previous reports.^38,39^

### Tissue Specificity

After cellular deconvolution, the proportion of cfDNA originating from the colon was not different between cancer patients with and without peritoneal metastases (Suppl Table 3). Cancer patients had a higher proportion of cfDNA from CD8 positive T-Cells (1.6% in PM^+^ samples, 1.5% in PM^−^ samples, and 0.9% in NC controls, Figure S 2 e). In contrast, NK cells contributed 6.0% in both PM^+^ and PM^−^ samples, which was lower than the 7.3% observed in NC controls (Supplementary Figure S 2 e).

### 5hmC Analysis

Differential 5hmC analysis between PM^+^ and PM^−^ samples identified 127 differentially 5hmC hydroxymethylated regions (DhMR) (p-value < 0.001 and |log2-fold change| > 0.2.) (Figure 2 a,b and Supplementary Table 4). Euclidean distance-based clustering (reinforced by PCA) with the same set of DhMRs was capable of separating PM^+^ samples from NC controls, with modest separation between PM^−^ samples to NC controls (Supplementary Figure S 3 a-h). We compared the 5hmC levels of the 127 DhMRs grouped separately into the 21 DhMRs enriched in PM^+^ and the 106 DhMRs enriched in PM^−^. In the PM^+^ enriched regions, PM^+^ samples showed significantly higher combined 5hmC levels than NC controls, whereas PM^−^ samples had lower levels of 5hmC than NC controls. In contrast, in the PM^−^ enriched regions, the opposite trends were observed (Supplementary Figure S 3 i).

**Figure 2:**
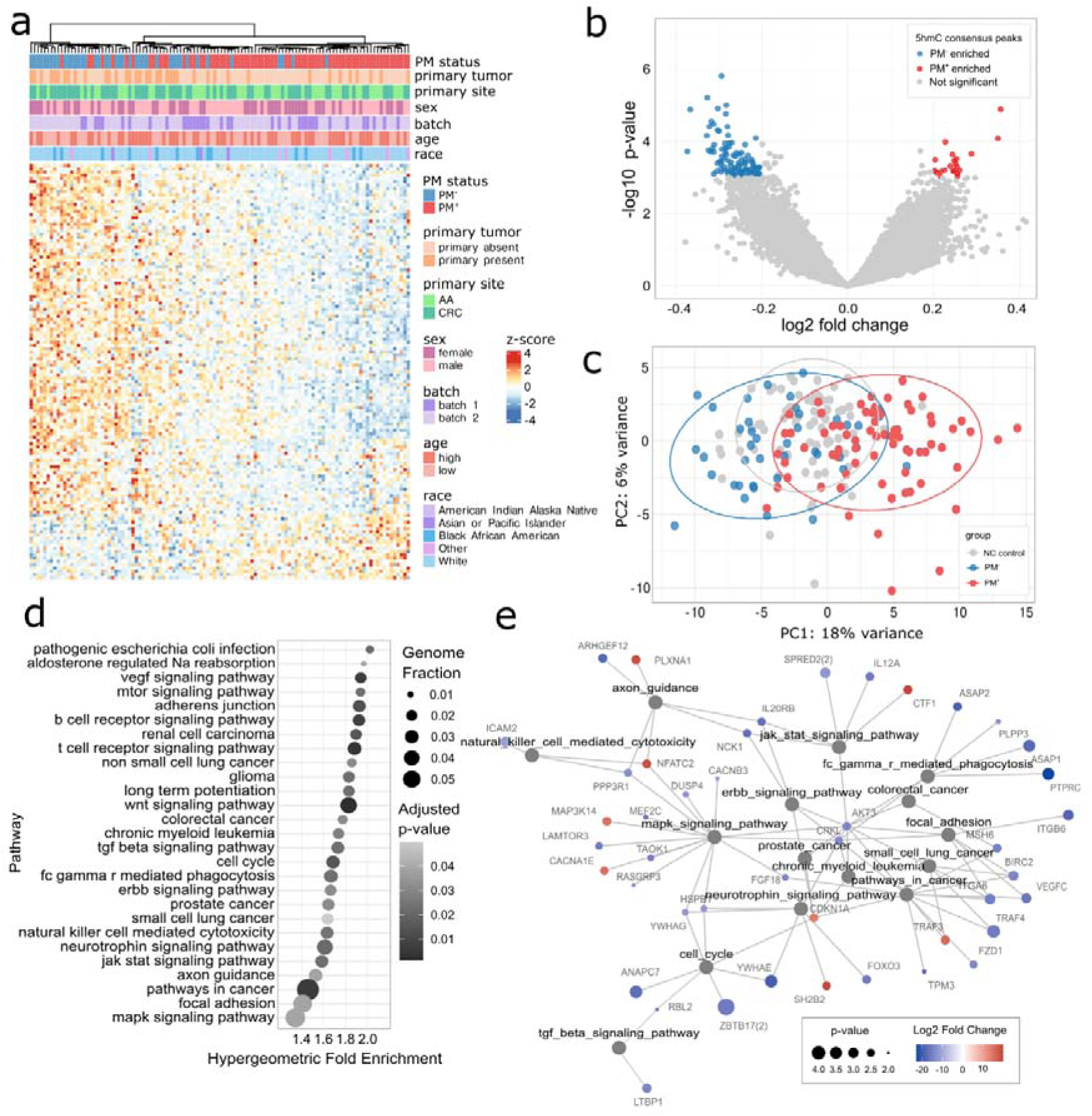
Functional analysis of differentially hydroxymethylated regions (DhMRs) between PM+ and PM-conditions. Differential 5hmC signatures are detectable between PM^+^ and PM^-^ samples in cfDNA obtained from patient peripheral blood samples. **a** Heatmap showing sample clustering by peritoneal metastasis status based on the z-score of 127 differentially hydroxymethylated regions (DhMRs) meeting statistical significance thresholds (Wald test p-value < 0.001 and |log2-fold change| > 0.2). Higher z-score indicates 5hmC enrichment in PM^+^ samples. Samples and DhMRs were clustered using Ward.D2 Euclidean distance metric. **b** Volcano plot showing the significance and magnitude of changes in DhMR 5hmC levels between PM^+^ and PM^-^ conditions. Points are color-coded based on statistical significance, and PM group in which they are enriched. **c** Principal component analysis (PCA) of 127 DhMRs from (a). PCA plot demonstrates separation between PM^+^ and PM^-^ samples, and greater overlap between PM^-^ samples and non-cancer (NC) controls relative to PM^+^ samples, suggesting a distinct epigenetic signature associated with the presence peritoneal metastases. Ellipses represent 90% confidence levels. **d** Dot plot depicting the top enriched KEGG pathways meeting a hypergeometric adjusted p-value cutoff of <0.05 based on the Genomic Regions Enrichment of Annotations Tool (GREAT) analysis of the top 2,000 DhMRs (greatest absolute value of the Wald statistic from DESeq2). Each dot represents a pathway. Dot size indicates the fraction of the entire genome that overlaps with regions associated with the given pathway. Color indicates the statistical significance of enrichment. **e** Network plot illustrating the relationships between the 15 most significantly enriched DhMRs (lowest hyper-adjusted p-value) and the top 200 DhMRs. Nodes represent individual DhMRs, and edges indicate connections between regions sharing similar KEGG pathway enrichments. Node labels represent either the annotated gene in which the DhMR is located, or the regulated target based on GeneHancer elite designation (see methods). The number in parentheses indicates that multiple DhMRs share the same annotated gene and/or regulated target. CRC: colorectal adenocarcinoma; AA: appendiceal adenocarcinoma; PC: principal component.

### Pathway analysis

27 pathways were significantly enriched including cancer related, immune related and signaling (including Wnt) pathways, (adjusted hypergeometric p-value <0.05) (Figure 2 d). Further, pathways related to cell migration and adhesion, which play a role in peritoneal dissemination, were identified such as “adherens junctions,” “axon guidance” and “focal adhesion”. Network analysis revealed that DhMRs associated with CRKL and AKT3 both contributed to the enrichment in the “focal adhesion,” “pathways in cancer,” “ErbB signaling pathway” and “chronic myeloid leukemia” pathways (Figure 2 e). These results suggest that enhanced invasive and/or proliferative potential, along with immune escape or exhaustion mechanisms regulated, at least in part, by epigenetic changes detectable in 5hmC levels in cfDNA, may be factors in the development of peritoneal metastasis.

### Regulatory region 5hmC levels in cfDNA are predictive of peritoneal metastasis

Of the 71,597 consensus hMR features available for training, we retained 5000 features after a Random Forest (RF) filter on outer-fold training sets prior to elastic net regularization modeling (Supplementary Figure S 4) (Figure 3 c).The mean outer fold AUC across 100 training runs was 0.827 (95% CI: 0.786 – 0.869) (Figure 3 a) with an average 92.4% sensitivity at 46.1% specificity, and 75.4% accuracy (Figure 3 b). As a negative control the same procedure was repeated with randomized PM status labels, which resulted in an, as expected, complete loss of predictive power in the outer folds to a mean AUC of 0.515 (95% CI: 0.450 – 0.580) (Figure 3 a) indicating that model performance is likely not the result of chance or overfitting.

**Figure 3:**
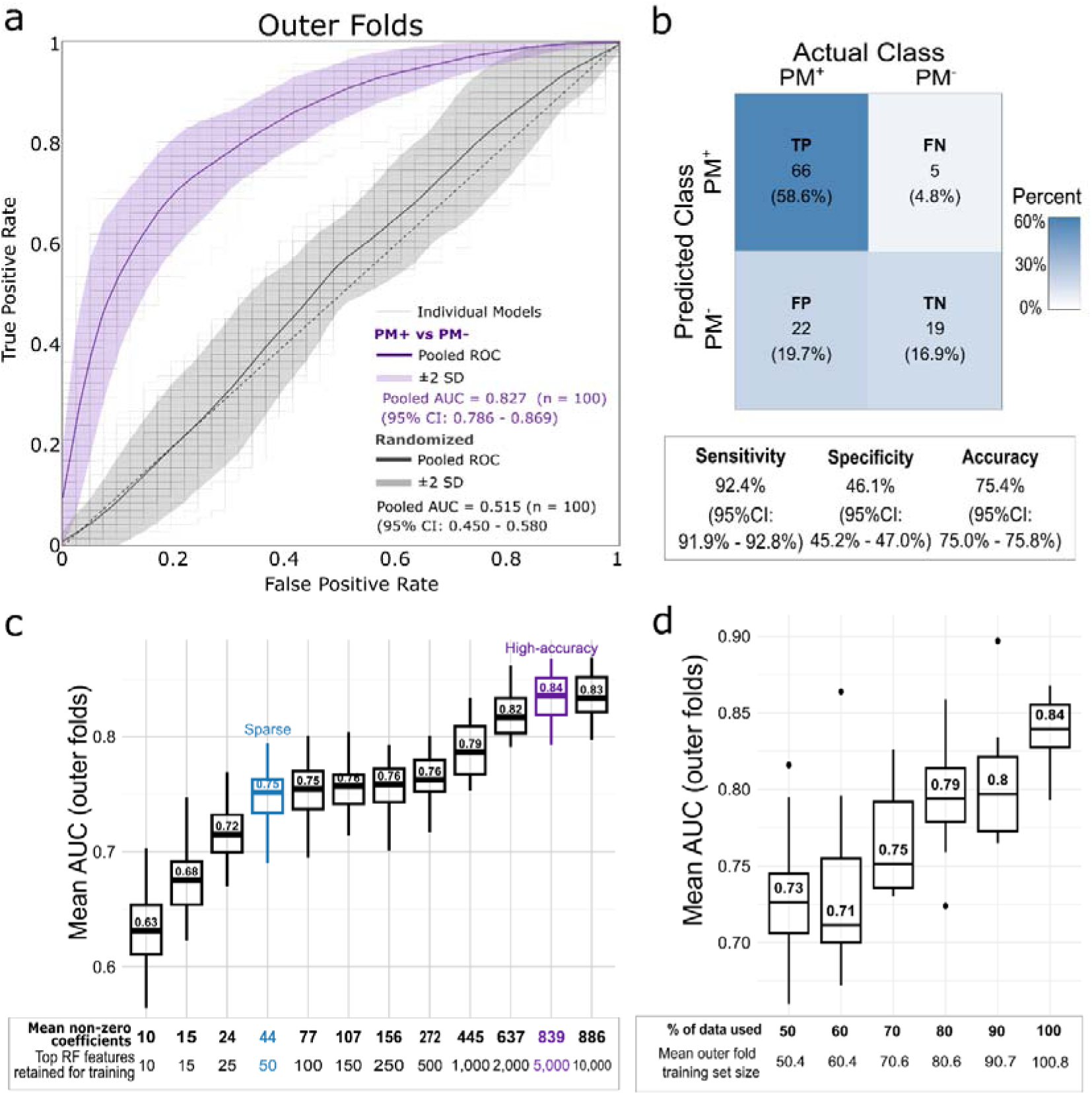
5hmC predictive model for Cancer and PM status. Consensus 5hmC peaks (n = 71,597) were used to train predictive models for peritoneal metastasis (PM) status. Briefly, 10-fold nested cross-validation (CV) was performed in combination with a random forest filter and elastic net regularization. This procedure was repeated 100 times varying the random seed. **a** Receiver operating characteristic (ROC) curves for nested CV outer folds. PM status models achieved a pooled outer-fold area under the curve (AUC) of 0.827 (95% CI 0.786-0.869) (n = 1,000) relative to 0.515 (95% CI 0.450-0.580) (n = 1,000) for the negative control with randomized PM status labels. **b** Pooled confusion matrix for all outer-fold test sets, for each run with a different random seed PM^+^ A sensitivity of 92.4% (95% CI 91.9-92.8%) was achieved, along with a specificity of 46.1% (95% CI 45.2-47.0%), and an accuracy of 75.4% (95% CI 75.0-75.8%). **c** Boxplot of mean outer-fold AUC relative to the number of features retained after Random Forest (RF) filtering, prior to elastic net regularization, and the mean number of non-zero coefficients after model training (n = 20 model runs varying random seed). Highlighted in blue is the RF parameter used for the sparse model predictive of PM status, and in purple the high-accuracy model predictive of PM status. **d** Boxplot showing mean outer fold AUC relative to the number of samples used for model training and evaluation showing an increase in model accuracy with increasing amounts of training data. (n = 20 model runs varying random seed).

### Model Assessment

Model accuracy increased with the amount of training data (Figure 3 d). Models trained with the highest accuracy parameters relied on a median of 851 hMRs with non-zero coefficients (Supplementary Figure S 7 e), with 27 hMRs occurring in greater than 99% of models (Supplementary Table 5). Among these 27 features, none flipped signs across runs (Figure 4 a), indicating that features were consistent in their association with either PM^+^ or PM^−^ status. Furthermore, hierarchical clustering of these features led to a strong separation of samples based on PM status (Figure 4 b). While the average predicted probability of PM stratified by primary site showed slight differences between CRC and AA samples (Figure 4 c), DeLong tests on the median ROC curves found that these differences were not statistically significant (p-value = 0.45) (Supplementary Figure S 5 a), suggesting that the models performed equally well at predicting PM status, regardless of the colorectal versus appendiceal origin of primary tumor. When comparing ROC curves stratified by sex, age, race, or presence of primary tumor, only the presence of the primary tumor at the time of plasma sampling showed a statistically significant difference in AUC (p-value = 0.046) (Supplementary Figure S 5 b-f), with higher performance in patients where the primary tumor was absent at the time of plasma sampling, likely a result of this group (primary tumor absent) being overrepresented in the PM^+^ class.

**Figure 4:**
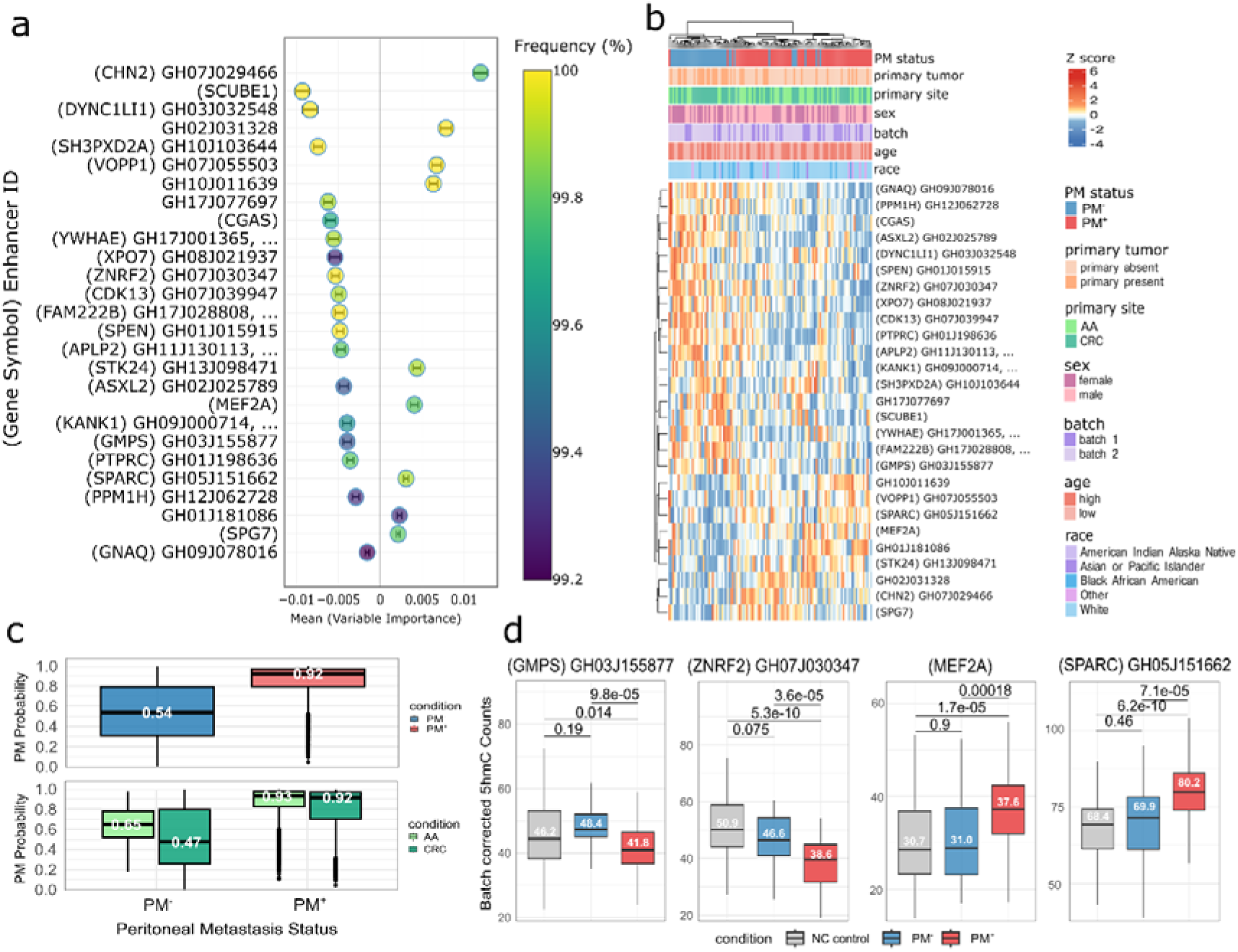
5hmC predictive performance across sub-groups, and coefficients are stable across runs. **a** Variable importance plot showing mean coefficients of DhMR features occurring in > 99% of models (n = 1,000) predictive of peritoneal metastasis (PM) status. Features show highly stable model coefficients, as shown by error bars representing pooled standard error of the mean (SEM). A coefficient > 0 indicates 5hmC enrichment in these genomic regions is associated with an increased probability of PM. **b** Heatmap showing sample clustering by PM status based on the z-score of 27 DhMRs from (a). Samples and DhMRs were clustered using the Ward.D2 Euclidean distance metric. **c** Model predictions of PM probability show strong separation between PM^+^ and PM^-^ samples (median outer-fold probability of 0.54 for PM^-^ samples relative to 0.92 for PM^+^) across all model runs (n = 11,200). Large sample size is due to each of the 112 samples being assigned a probability prediction for each random seed (n = 100). **d** Average 5hmC batch corrected counts for DhMRs associated with select genes based on presence in gene body, or GeneHancer Elite designation for enhancer-gene association. All are examples demonstrating statistically significant differences (adjusted p-values < 1e-3, Wilcoxon test with adjusted by the Benjamini-Hochberg method) in DhMR 5hmC levels in PM^+^ relative to PM^-^ and/or non-cancer controls, where associated genes were found to have statistically significant (FDR <0.05) survival differences based on high/low expression in TCGA database. CRC: colorectal adenocarcinoma; AA: appendiceal adenocarcinoma; NC: non-cancer; PM: peritoneal metastasis.

### Sparse models predictive of PM status show similar properties to high accuracy models

We evaluated the performance of sparse elastic net regularization models trained after retaining 50 features following RF filtering, rather than the 5,000 used in the high-accuracy models (Figure 3 c). The mean outer fold AUC was 0.749 (95% CI: 0.691 – 0.808) (Supplementary Figure S 6 a) with an average 86.3% sensitivity at 45.6% specificity, and 71.4% accuracy (Supplementary Figure S 6 a-b). Models trained with this sparse parameter set relied on a median of 44 hMRs with non-zero coefficients,12 of which also occurred in >99% of high-accuracy PM models 17 hMRs occurring in greater than 50% of models (Supplementary Figure S 6 c, e, Supplementary Table 6). Similar to the high-accuracy models, sparse model performance was limited by training data availability (Supplementary Figure S 6 d).

A previous report found that a sparse model trained with 5hmC levels in cfDNA was predictive of CRC status relative to NC controls.^40^ As a positive control to confirm that a higher accuracy of a sparse model (albeit of cancer status, and not PM status) is achievable with our small dataset, we used our data to replicate this finding by training a set of models to predict cancer status (CRC or AA). Consistent with previous reports, on average, our models were able to predict the cancer status of patients relative to NC controls with an outer-fold average AUC of 0.925 (95% CI: 0.904 – 0.946) (Supplementary Figure S 7 a) with an average 76.1% sensitivity at 90.6% specificity and 84.9% accuracy (Supplementary Figure S 7 c). The median sparse model predictive of cancer status relied on 45 hMR features (Supplementary Figure S 7 d, e, Supplementary Table 7) which were able to effectively separate cancer from non-cancer samples by hierarchical clustering (Supplementary Figure S 7 b).

### hMRs predictive of PM status are associated with known cancer metastasis-relevant genes

Of the 27 hMRs occurring in >99% of high-accuracy models predictive of PM status, 23 hMRs map to known enhancer regulatory regions, two are located in intronic non-enhancer regions, and two are located in 3 or 5’ UTRs (Supplementary Table 5). Upon associating all enhancer hMRs to high-confidence regulated target genes, based on the GeneHancer elite designation,^41^ we obtained a list of 28 genes (15 of which are based on the GeneHancer gene-enhancer associations, and 13 of which are based on the hMR being located within the associated gene body). Of these 28 hMR-associated genes, 14 were found to have their mRNA gene-chip expression levels significantly associated with an increase or decrease in relapse free survival (RFS) amongst 1,167 CRC patients in The Cancer Genome Atlas (TCGA) database accessed via KM plotter (p-value ≤ 0.001, Supplementary Table 8).^42^ We performed a permutation test with 10,000 iterations to compare the median p-value of RFS differences in CRC patients for this set of hMR-associated genes (median p-value = 1.15e-03, n=28), with that of set of over 1,000 genes drawn at random (median p-value = 9.00e-3, n=1,061). hMR-associated genes were found to have lower median p-values, indicating a statistically significant enrichment for associations with RFS differences in CRC patients (p-value = 4.68e-2) (Supplementary Figure S 8a). Furthermore, of these 28 genes, 23 were found to have established links to CRC in the literature (Supplementary Table 9).

For example, the hMR located in enhancer GH05J151662 in the 3’ UTR of Secreted Protein Acidic And Cysteine Rich (*SPARC)* gene is significantly enriched in PM^+^ patients (p-value < 1e-4) relative to PM^−^ patients (Figure 4 d), and relative to NC controls patients (p-value < 1e-9). High mRNA expression of the SPARC gene is associated with a statistically significant decrease (p-value < 1e-4) in 5-year RFS in CRC patients in the TCGA database (Supplementary Figure S 8 b). In contrast, the hMR overlapping enhancer GH07J030347 in an intron of Zinc and Ring Finger 2 (*ZNRF2*) is significantly enriched in both PM^−^ patients (p-value < 1e-4) and NC controls (p-value < 1e-9) relative to PM^+^ patients (Figure 4 d). High mRNA expression of the *ZNRF2* gene is associated with an increase in 5-year RFS (p-value < 1e-4) in CRC patients in the TCGA database (Supplementary Figure S 8 c).

## Discussion

To the best of our knowledge, our findings demonstrate proof of concept that 5hmC signatures detectable in cfDNA can effectively distinguish PM^+^ patients from PM^-^ patients at sensitivity levels higher than currently available detection modalities albeit with modest specificity.^43–46^ Our results align with previous studies that have identified 5hmC-modified genes and genomic regions as putative biomarkers in cfDNA for distinguishing cancer patients from non-cancer individuals^16–21^. This is the first study to extend this approach to patients with PM.

The performance of our test (AUC 0.83) in distinguishing PM^+^ from PM^-^ patients, and in identifying cancer patients (CRC or AA) from NC controls (AUC 0.94), is likely due to our enabling 5hmC-Seal chemical labeling technology that allowed us to measure genome-wide 5hmC levels from nanograms of cfDNA, which appears to be superior to previously reported cell free and circulating tumor DNA technologies. Furthermore, the increasing accuracy of the models with larger training datasets suggests that further performance gains are likely to be achieved with an expanded sample size. It is presently unknown whether the performance difference between patients with and without primary tumor at the time of sample is a result of actual differences in 5hmC signatures, or a result of our imbalanced dataset. Future studies with larger datasets should consider these groups separately to clarify this observation.

Pathway analysis of DhMRs revealed four main clusters of pathways epigenetically dysregulated in PM related to cancer, the immune system, cell signaling and migration pathways in patients with peritoneal metastasis, suggesting two major mechanisms by which epigenetic dysregulation might contribute to PM. Firstly, enrichments of pathways involved in cell migration, and adhesion suggests that dysregulation in these pathways could enhance the motility and invasive capacity of tumor cells. Specifically, DhMRs associated with genes like *PLXNA1*, *ARHGEF12*, *AKT3, ITGA6, BIRC2,* and *CRKL* (Figure 2 d) is consistent with a model in which epigenetic regulation of these genes or their pathways may promote the detachment of tumor cells from the primary site, increasing cancer cell survival in the peritoneal environment, or enhancing their invasive capabilities, leading to the establishment of metastatic lesions. Secondly, the epigenetic dysregulation of immune-related pathways suggests that evasion of immune surveillance or other immune system interference might play a role in metastasis to the peritoneum. Specifically, epigenetic dysregulation of DhMRs associated with genes such as *ICAM2*, *NFATC2*, *PPP3R1*, *PTPRC*, and *LTBP1* (Figure 2 d) might impair the immune system’s ability to recognize and eliminate tumor cells. Immune evasion or immune cell exhaustion could contribute to metastasis by allowing tumor cells to survive and proliferate in the peritoneum.

Many genes associated with the 5hmC-enriched regions relied upon by our PM-predictive models have known roles in cancer metastasis. For instance the *SPARC* gene, whose 3’ UTR enhancer was significantly enriched for 5hmC in PM^+^ patients, was found in previous studies to be differentially expressed (4.6 fold change, 0.02 FDR) in CRC tissues with PM relative to those without,^47^ as well as in CRC cell lines able to develop high numbers of peritoneal lesions in mice (2.1 fold change, 2.91E-14 FDR).^48^ The *ZNRF2* gene on the other hand, which was found in a previous study to have lower expression in CRC cell lines able to develop high numbers of peritoneal lesions in mice (−1.6 fold change, 2.91E-14 FDR),^48^ had higher 5hmC levels in its intronic enhancer GH07J030347 in PM^-^ patients, consistent with increased expression and a potential tumor suppressor role in PM^-^ patients. Taken together, these results show that many of the genes associated with hMRs predictive of PM status, have known links with CRC metastasis. Unfortunately, AA is a rarer and less well studied disease relative to CRC. As a result, similar analyses of published literature of AA are not presently available. These pathways and genes warrant further mechanistic exploration and validation in future studies.

The ability to detect PM from a minimally invasive blood-based assay has significant clinical implications. Patients with peritoneal metastases if untreated die within weeks to months. Among those treated with systemic therapy, the median survival is a modest 14-18 months with no long-term survivors. Curative intent surgery (CRS) in select patients with low burden of disease has demonstrated favorable survival with 90% 5 year survival and over 40-60% cure rate.^49–52^ It is therefore conceivable that routine application of such a test for high-risk CRC and AA patients could identify patients with low to moderate burden of disease that would benefit from surgery and/or could participate in novel clinical trials. A low burden of disease also correlates with lower morbidity from surgery and the ability to apply minimally invasive surgical techniques that would reduce morbidity for patients.

In addition to identifying patients who benefit from curative intent therapy, this test could also de-escalate therapy for patients, particularly those that undergo routine diagnostic laparoscopy for detection of peritoneal disease, or those that receive adjuvant systemic chemotherapy for high-risk colon cancer. Similar designs have been proposed with the use of ctDNA in de-escalation of therapy with promising results.

Biomarkers for diagnosis of peritoneal metastases if validated for clinical benefit, could be deployed at diagnosis of cancer, used for surveillance of treated high risk (T4 colon/appendix cancer, perforated cancers) and metastatic cancers in conjunction with imaging and other plasma tumor markers. However, given the lower specificity of our signature, we recognize the need to deploy the test appropriately to obviate patient anxiety with false positive results.

Our study has several limitations. First, to our knowledge, while this is one of the largest biomarker discovery cohorts of patients with PM reported in the literature to date, the sample size remains relatively small which prevented a separate analysis of CRC and AA patients and limited our ability to draw specific conclusions that might have distinguished these diseases. Larger independent cohorts are necessary to separately validate our findings in CRC and AA, to identify a specific panel of genomic regions for use in a future clinical assay, and to extend our results to other cancers known to metastasize to the peritoneum such as ovarian, gastric or pancreatic cancer. Finally, longitudinal studies are needed to assess the temporal and causative relationship between the 5hmC alterations identified in this study, and PM development.

Future studies integrating 5hmC signatures with other biomarkers, such as circulating tumor DNA, protein biomarkers, or other epigenetic marks, could further improve the diagnostic accuracy of a minimally invasive blood-based assay for PM. Future work exploring the biological mechanisms underlying the 5hmC changes could also uncover new therapeutic targets for PM. Finally, investigating the utility of 5hmC signatures to monitor patient response to treatment and detect disease recurrence could also be valuable. Given the minimally invasive nature of plasma-derived cfDNA collection, serial sampling is feasible and could offer real-time insights into disease dynamics.

Our study demonstrates that 5hmC signatures from plasma-derived cfDNA could serve as novel biomarkers for detecting PM in patients with CRC or AA. While larger studies and additional validation cohorts are necessary, our findings open avenues for improving the management of PM through minimally invasive blood sampling and a better understanding of the molecular mechanisms contributing to PM.

## Acknowledgements and funding

We would like to acknowledge Alana Beadell for her assistance with 5hmC sample preparation, Varun Bansal and Leah Ulrich for helping coordinate research meetings and accruing data and Dr. Oliver Eng for assisting with project administration. We would also like to acknowledge the support of various large language model (LLM) tools such as ChatGPT by OpenAI, Claude by Anthropic and Copilot by GitHub for coding assistance.

We would like to acknowledge the Irving Harris Foundation, The Kevin Brown Family Foundation and the Gastrointestinal Research Foundation that graciously supported this study.

## Supplementary methods

### Blood Sample and Processing

Around five milliliters of peripheral blood were collected from the patients in EDTA containing purple top tubes. The average cfDNA extracted per sample was 9.7 ng (IQR: 9.7-10.0) (Supplementary Figure S 1). Briefly, within 6 hours, plasma was extracted using the Quick-cfDNA Serum & Plasma Kit (ZYMO) by centrifuging twice at 1350×g for 12 min at 4 °C and 13,500×g for 12 min at 4 °C. Then, the plasma samples were immediately stored at – 80 °C until further use. Then, plasma samples were processed for cfDNA extraction using the QIAamp Circulating Nucleic Acid kit (Qiagen, Valencia, CA), as per the manufacturer guidance. The fragment size of all the cfDNA samples were verified by nucleic acid electrophoresis before library preparation.

### 5hmC library construction and high-throughput sequencing

5hmC libraries for all samples were constructed with high-efficiency 5hmC-Seal technology.^21^ According to the requirements of next-generation sequencing, the cfDNA extracted from plasma was end-repaired, 3′-adenylated using the KAPA Hyper Prep Kit (KAPA Biosystems) and then ligated with the Illumina compatible adapters. The ligated cfDNA was added in a glycosylation reaction in 25 μL solution containing 50 mM HEPES buffer (pH 8.0), 25 mM MgCl2, 100 μM UDP-6-N3-Glc, and 1 μM β-glucosyltransferase (NEB) for 2 h at 37 °C. Next, the cfDNA was purified using DNA Clean & Concentrator Kit (ZYMO). The purified DNA was incubated with 1 μL of DBCO-PEG4-biotin (Click Chemistry Tools, 4.5 mM stock in DMSO) for 2 h at 37 °C. Similarly, the DNA was purified using the DNA Clean & Concentrator Kit (ZYMO). Meantime, 2.5 μL streptavidin beads (Life Technologies) in 1 × buffer (5 mM Tris pH 7.5, 0.5 mM EDTA, 1 M NaCl, and 0.2% Tween20) were added directly to the reaction for 30 minutes at room temperature. Finally, the beads were subsequently washed eight times for five minutes with buffer 1–4. All binding and washing steps were performed at room temperature with gentle rotation. Then, the beads were resuspended in RNase-free water and amplified with 14–16 cycles of PCR amplification. According to the manufacturer’s instructions, the PCR products were purified using AMPure XP beads (Beckman). The concentration of libraries was measured with a Qubit 3.0 fluorometer (Life Technologies). Paired-end 50-bp high-throughput sequencing was performed on the Novaseq 6000.

### Reads trimming, quality control and mapping

All reads from each samples’ two paired-end “.fastq” files had sequencing adapters trimmed using Trim Galore^53^ (version 0.6.7) with following parameters: “--paired” for paired-end mode and “--length 15” to discard reads shorter than 15bp after trimming.

Sequencing quality control on raw “.fastq” files was performed using FastQC (version 0.11.9). Results for each sample were collated and analyzed using MultiQC (version 1.21). Samples with at least one of two paired-end files with fewer than 10M unique reads were excluded from all subsequent analysis due to library preparation and/or sequencing failure. This resulted in the exclusion of 7/192 (3.6%) of samples (Supplementary Figure S 1).

Reads from both trimmed paired-end “.fq” files for each sample were then mapped to the human genome assembly GRCh37 (hg19) using Bowtie 2 (version 2.4.4) with the following parameters: “-N 1” allowing up to 1 mismatch in seed alignment. Resulting .bam files were then sorted and indexed using Samtools (version 1.16.1).

For all unspecified parameters, default values were used.

### 5hmC consensus peak set, and read counts determination

5hmC peaks were called on each sample .bam file using MACS3 version (3.0.1) with the following parameters: “--nolambda” to account for the absence of a negative control for the 5hmC-Seal sequencing method, “-f BAMPE” specifying paired-end format of input .bam file, “-p 1e-5” to set a permissive cutoff and “-g hs” to specify the genome size.

All samples’ resulting “.xls” peak files were then combined into a merged peak-set using DiffBind (version 3.12.0) “dba” and “dba.peakset” functions with the following parameters respectively: “doBlacklist=DBA_BLACKLIST_HG19” to exclude peaks from ENCODE hg19 blacklist regions, “minOverlap=0.1” to select peaks that occurred in at least 10% of samples.

Raw 5hmC-associated read counts for each sample in merged peak-set regions were then determined using featureCounts (version 2.0.3) with the following parameters: “-p” to indicate paired-end mode, “--countReadPairs” to ensure that read pairs rather than individual reads were counted, “-Q 20” to require a minimum mapping quality score of 20, “-c” to exclude reads that mapped to multiple locations in the genome, and “--ignoreDup” to ignore duplicate reads potentially introduced during PCR amplification.

For all unspecified parameters, default values were used.

The resulting merged peak set containing 166,945 hMRs was further filtered into a consensus peak containing 71,597 by excluding peaks occurring on chromosomes X and Y, peaks greater than 5,000 bp in length, and peaks that had fewer than 15 counts in > 10% of samples (Supplementary Figure S 1 c, d).

### Region annotation and differential 5hmC enrichment analysis

All regions were annotated using annotatr (version 1.28.0) built-in hg19 annotation files ‘hg19_basicgenes’, or a user-provided annotation file ‘GeneHancer_hg19_v5.20’ provided by LifeMap Sciences. Enhancer – Gene association scores were only used for those associations annotated as “elite” where the association is based on at least two independent evidence sources.

5hmC metagene plot was constructed using deepTools (version 3.5.5) “computeMatrix” function, run on all genes in the hg19 genome assembly, with the following parameters: “--skipZeros” to skip samples with no reads, “-b 3000”, “-a 3000” and “--regionBodyLength 3000” in order to tabulate reads from 3000bp before and after all genes.

Differentially enriched 5hmC regions (DHRs) between conditions were determined using DESeq2 (version 1.42.1). A combined p-value and log fold change cutoff or an adjusted p-value cutoff was then applied depending on the use case (values specified in figures.).

All heatmaps were generated with ComplexHeatmap (version 2.18.0), with all rows and columns clustered with a Ward.D2 Euclidean distance measure.

### Cellular deconvolution analysis

To construct a comprehensive reference for deconvolution, we analyzed the 5hmC-Seal profiles from 20 different normal tissue and cell types from previous work,^22,54^ and selected the top 100 genes for each tissue and cell type. Reads per kilobase million (RPKM) values were calculated for both reference profiles and sample cfDNA profiles before deconvolution using CIBERSORT.^55^

### Pathway analysis

The 2,000 DhMRs with the greatest absolute value Wald statistic from DESeq2 were analyzed using rGREAT (version 2.5.7),^34^ to identify enriched gene ontology terms in the msigdb:C2:CP:KEGG database with a minimum geneset size of 15, using the default algorithm for region-gene assignments. Pathways with a hypergeometric adjusted p-value <0.05 were plotted in a network diagram along with their associated DhMRs, using ggraph (version 2.2.1), and in a dot plot using ggplot2 (version 3.5.1)

### Predictive model training and evaluation

All ∼71,597 hMRs were considered candidate features. An elastic net regularization model was then trained and evaluated using nested CV with 10 inner folds and 10 outer folds stratifying samples to maintain similar proportions of both sample PM status and sample primary site (e.g. CRC or AA) across folds. Class weighting was applied considering only PM status. Model training was performed using NestedCV (version 0.7.8) function “nestcv.glmnet”,^35^ with the following parameters: “standardize = TRUE” indicating to cv.glmnet to transform all hMRs raw read counts to have a mean of 0 and standard deviation of 1 prior to training, “alphaSet = seq(0.3,1, by = 0.1)” in order to limit the search space for the elastic net mixing parameter α, “min_1se = 1” to select the most regularized model such that the cross-validated error is within one standard error of the minimum,^56^ “finalCV = TRUE “ indicating to train a final model on the entire dataset, “filterFun = ranger_filter” indicating the use of ranger’s (version 0.16.0) Random Forest Filter with the following filter options: “num.trees = 10000”, “max.depth = 30”, “min.node.size = 20”, “nfilter = 150”, “mtry = 0.2*number_of_regions”. Grid search was performed to identify optimal RF filter parameters (data not shown). Top 50 (sparse) or 5,000 (high-accuracy) features resulting from the RF filter were then used for elastic net regularization model training. Nested CV was repeated 100 times varying the random seed. The occurrence frequency and coefficient for all hMRs in each run’s outer fold models were recorded (procedure illustrated in Supplementary Figure S 4). For all unspecified parameters, default values were used.

## Supplementary Figures

**Supplementary Figure S1:**
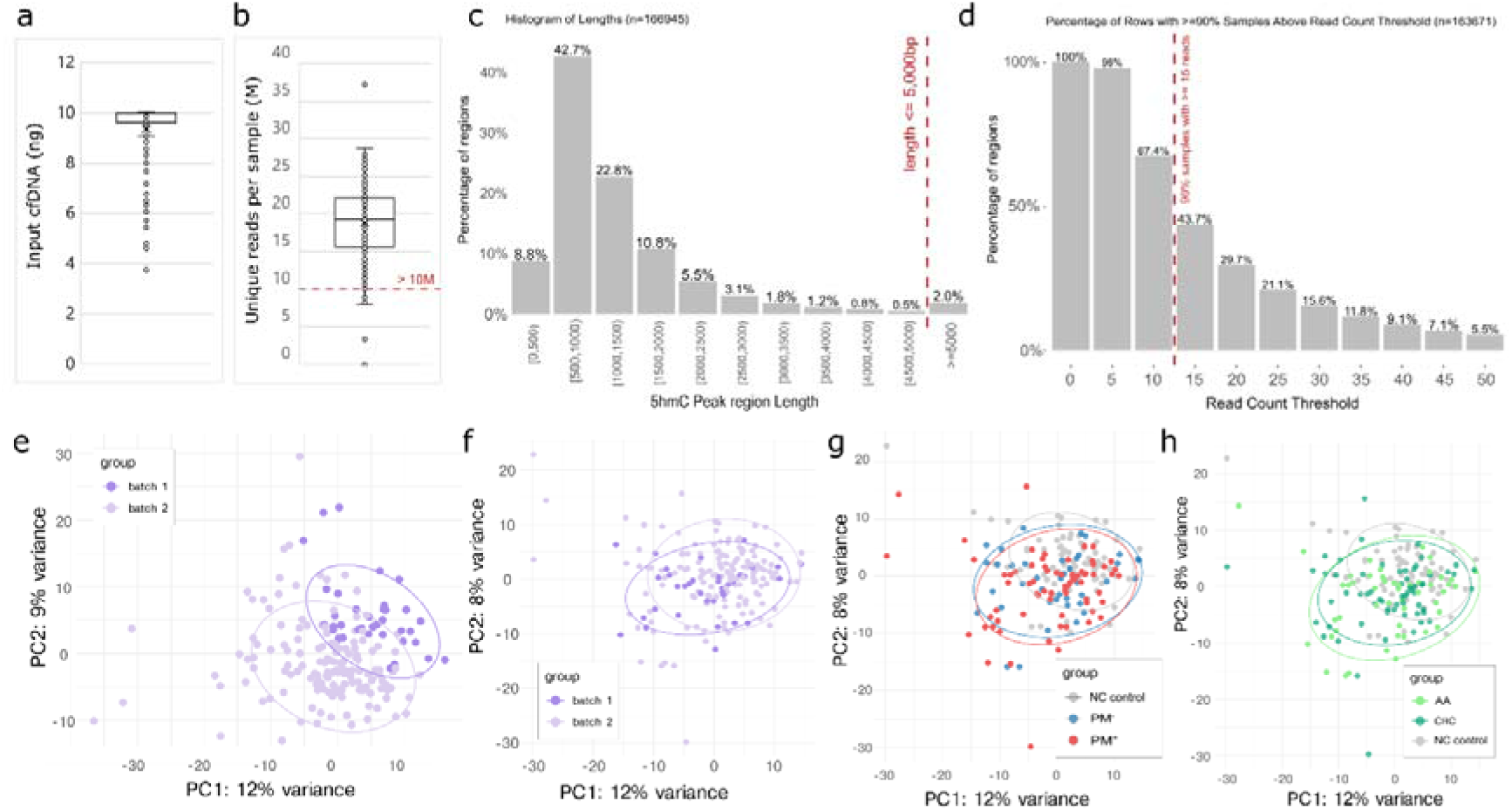
Input cfDNA, sequencing and genome-wide PCA quality control. **a** Box plot of input cell-free DNA (cfDNA) obtained per sample. An average of 9.4 ng of cfDNA was successfully extracted from each sample and used as input into the 5hmC-Seal protocol. **b** Box plot of minimum unique reads per paired end file. Samples with fewer than 10M unique reads were excluded from downstream analyses. **c** Bar plot of the distribution of hydroxymethylated region (hMR) lengths from merged peak set (n = 166,945). hMRs greater than 5,000bp in length were excluded from downstream analyses. **d** Bar plot showing percent of hMRs from merged peak set with ≥ 90% of samples with increasingly high read count thresholds (n = 163,671). Only hMRs with greater ≥ 15 reads in ≥90% of samples were retained in the consensus peak set, leaving 71,597 consensus hMRs for downstream analyses. **e** Genome-wide principal component analysis (PCA) plots based on raw cfDNA 5hmC counts across 71,597 5hmC consensus peaks showing batch effects between samples sequenced in batch 1 and 2. Ellipses represent 90% confidence level. **f** Same plot as (e) except with 5hmC counts adjusted using ComBat-seq to account for sample batch, showing elimination of batch effect. **g, h** Same plot as (f) with samples colored by peritoneal metastasis (PM) status and primary tumor site respectively, suggesting no genome-wide detectable differences between the plotted conditions, and the successful elimination of batch differences by the ComBat-seq normalization step. cfDNA: cell-free DNA; PC: principal component.

**Supplementary Figure S2:**
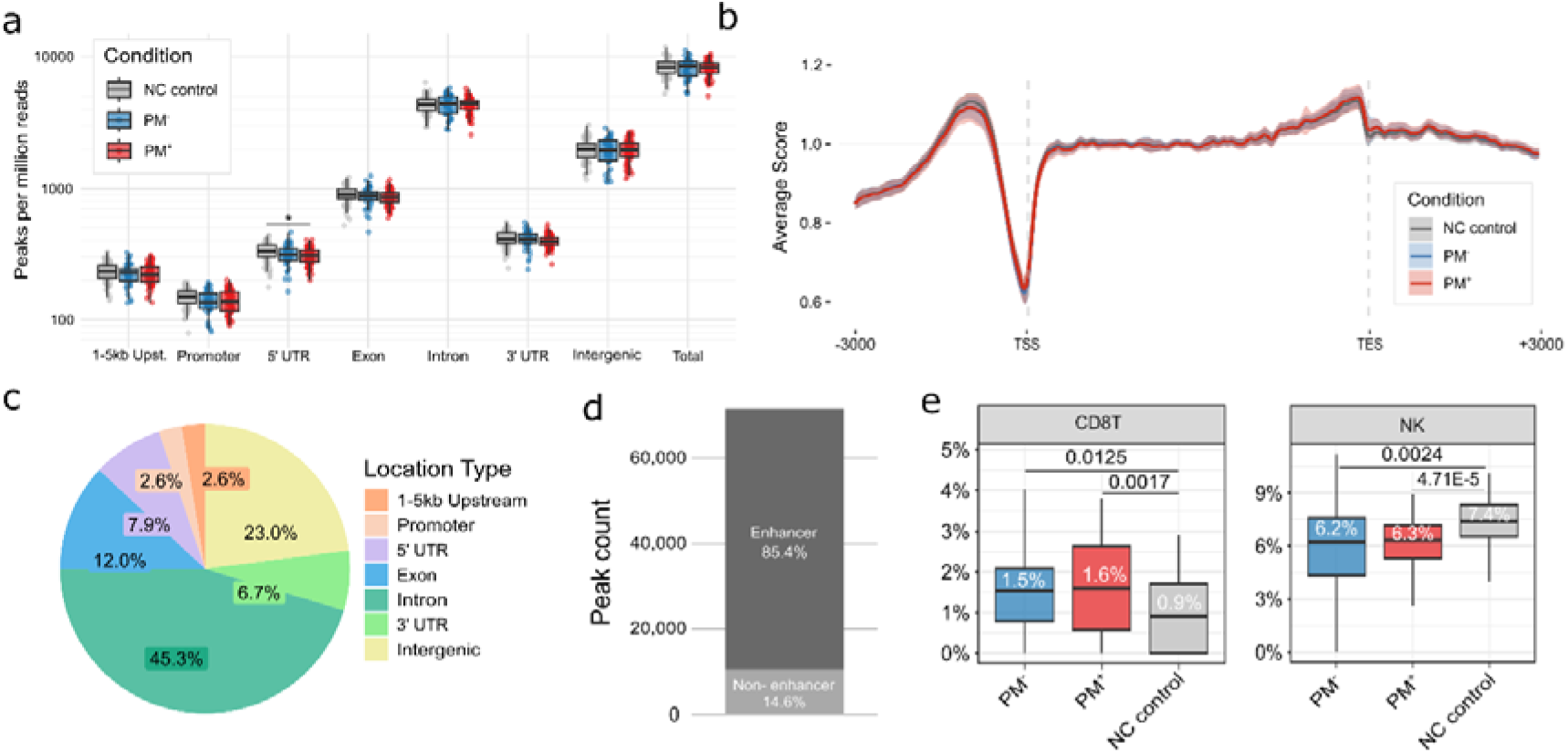
Genome-wide distribution of 5hmC consensus peaks. **a** Boxplots comparing normalized 5hmC peak density across different genomic feature types (1-5kb upstream, promoter, 5’ UTR, exon, intron, 3’ UTR, intergenic, and total) between conditions. Limited genome-wide changes in 5hmC peak density are detectable between non-cancer controls, PM^+^ and PM^-^ samples. Only 5’ UTR regions show a statistically significant decrease in peak enrichment between non-cancer controls and PM^+^ samples (* = p-value < 0.05, pairwise t-tests with Holm’s correction for multiple comparisons). **b** Metagene plot of normalized 5hmC enrichment score within ±3,000 bp of transcription start site (TSS) and transcription end site (TES) of 19,100 genes in the human genome, grouped by sample condition. Shaded regions represent one standard deviation from the mean. All conditions show characteristic 5hmC enrichment in promoters and gene bodies, and depletion surrounding the TSS, with no minimal differences between conditions. **c** Pie chart showing the distribution of consensus 5hmC peaks (n = 71,597) across genomic region types. A large majority of 5hmC peaks (71.8%) mapped to gene bodies. **d** Stacked bar chart shows the vast majority (85.4%) of all consensus peaks (n = 71,597) mapped to known enhancer regions. **e** Cell type deconvolution based on genome-wide 5hmC read counts shows that the cell-free DNA contribution of CD8^+^ T Cells is greater in PM^+^ and PM^-^ samples relative to non-cancer controls whereas natural killer (NK) cell contribution is greater in PM^-^. Pairwise Wilcoxon rank sum test was used for statistical significance. Upst.: upstream; UTR: untranslated region; NC: non-cancer.

**Supplementary Figure S3:**
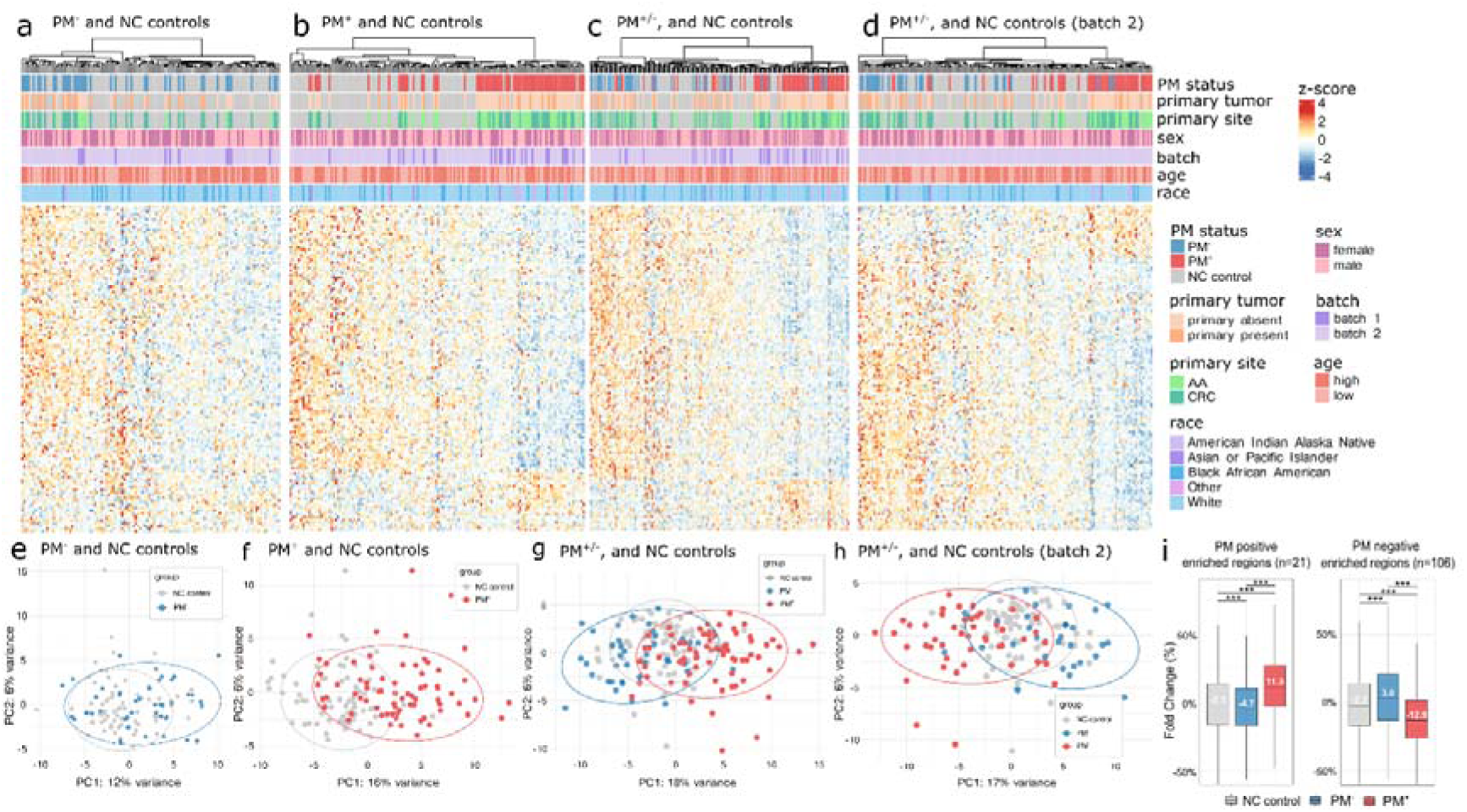
Heatmaps for each PM status relative to non-cancer controls for PM DhMRs. **a–d** Heatmaps display sample clustering by peritoneal metastasis status based on the z-score of 127 differentially hydroxymethylated regions (DhMRs) identified in cell-free DNA (cfDNA) extracted from peripheral blood. Samples and DhMRs were clustered using Ward.D2 Euclidean distance metric. Panels (**a**), (**b**), (**c**) show respectively: PM^-^ samples, PM^+^ samples and both combined with respect to non-cancer controls. Panel (**d**) shows both combined with respect to non-cancer (NC) controls from sequencing batch 2. **e–h** Principal component analysis (PCA) of 127 DhMRs corresponding to corresponding heatmaps from (**a**–**d**). Ellipses represent 90% confidence levels. PCA plot demonstrates separation between PM^+^ and PM^-^ cases, and a greater overlap between PM^-^ and NC controls suggesting a distinct epigenetic signature associated with the presence of peritoneal metastases. Panel (**g**) is repeated from Figure 2 for clarity. **i** Boxplots of fold-change percentage in 106 and 21 DhMRs identified in (**a**) enriched in PM^-^ and PM^+^ samples respectively, relative to NC controls. Medians are displayed in white within each box. Difference in means was assessed using ANOVA followed by Tukey’s Honestly Significant Difference post-hoc test for pairwise comparisons. Statistical significance indicated as follows: p-value < 0.05 (*), <0.01 (**) and <0.001 (***). The level of 5hmC in NC controls is much closer to that of PM^-^ samples, than PM^+^, suggesting these DhMRs are specific to PM status. CRC: colorectal adenocarcinoma; AA: appendiceal adenocarcinoma; NC: non-cancer; PM: peritoneal metastasis; PC: principal component.

**Supplementary Figure S4:**
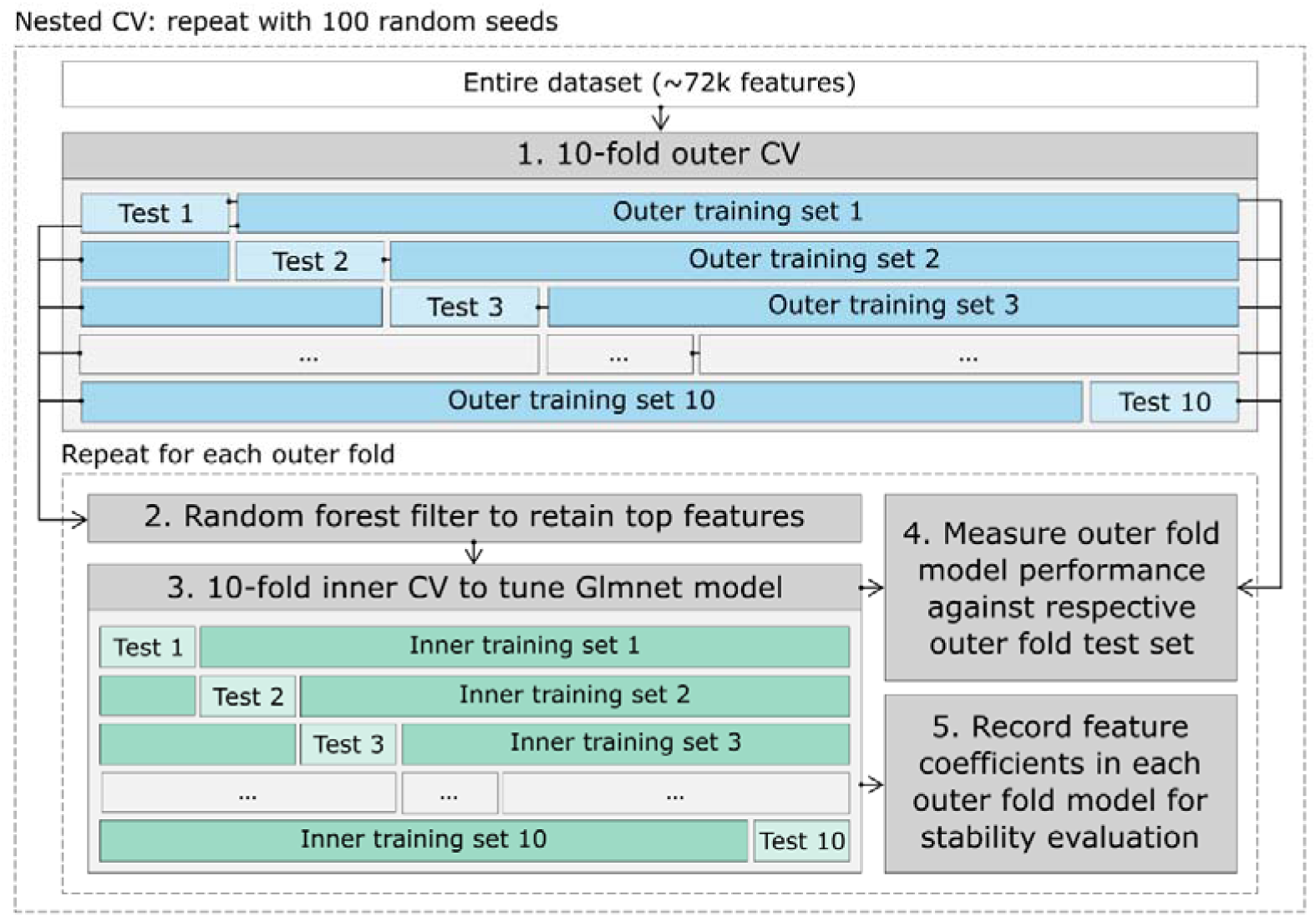
Model training procedure flow chart. Detailed nested cross-validation (CV) procedure used for training and evaluating performance of predictive models. Nested CV was selected as an optimal approach given the small sample size available in this study. Briefly (1) 10-fold outer cross validation was performed on all features (n = 71,597 consensus hMRs) whereby outer training sets were used for subsequent feature filtering and training, and outer test sets were held out as a validation set. (2) A random forest filter was used to select the top features for subsequent elastic net regularization. (3) 10-fold inner cross-validation was used to optimize elastic net alpha and lambda parameters. (4) Optimal model parameters were then used to train a model on all samples in the outer training set, and performance was evaluated against the outer test set which was held out for validation. This procedure is repeated for each of the ten training – test splits, allowing each sample to occur in an outer test set once. (5) Feature coefficients and occurrence frequency in each outer-fold model were recorded for further analysis. The entire procedure was repeated 100 times varying random seed.

**Supplementary Figure S5:**
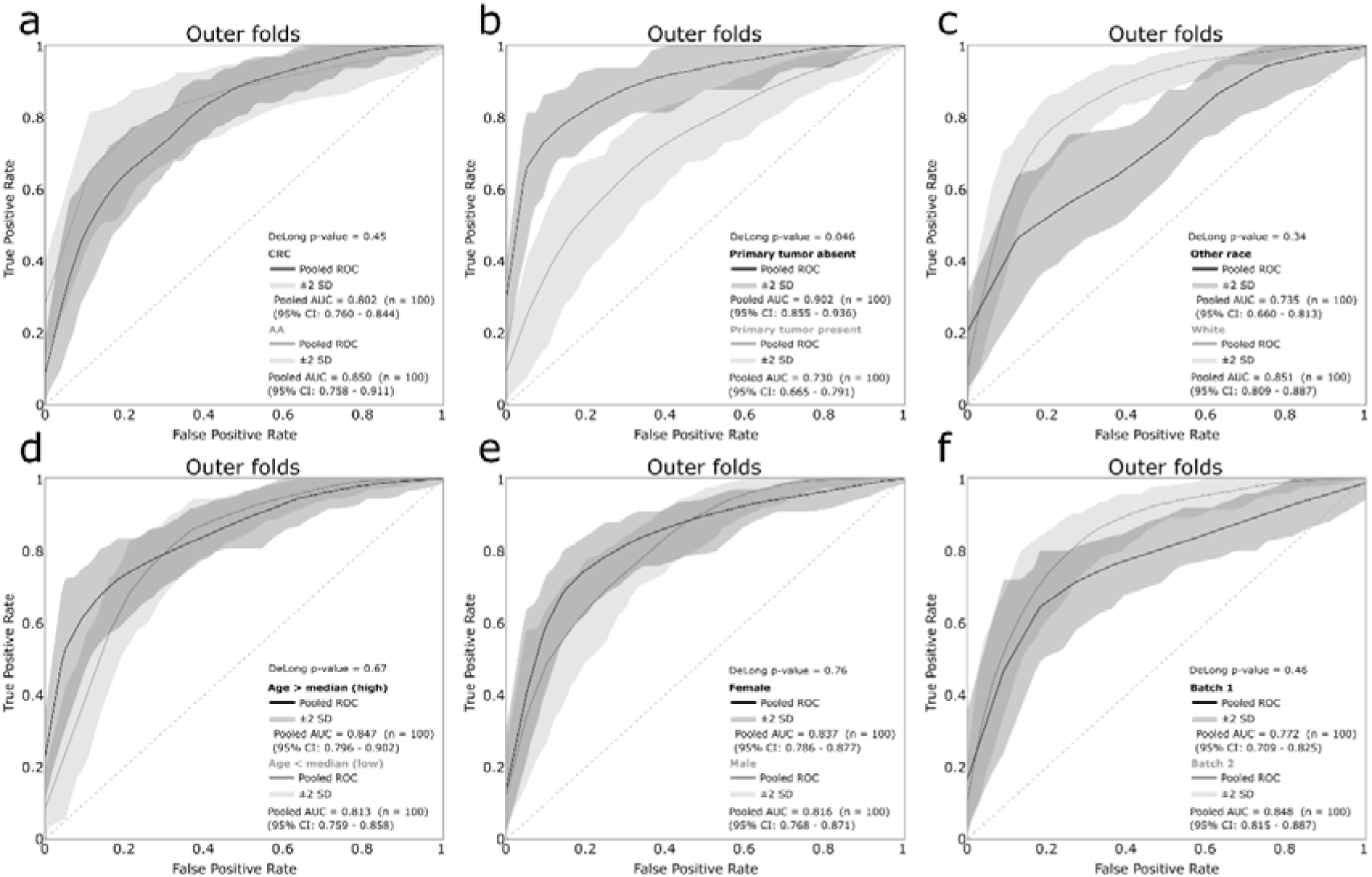
PM model performance stratified by sub-group. **a–f** Receiver operating characteristic (ROC) curves for outer folds from of elastic net regularization models of PM status, stratified by primary site, presence of primary tumor at time of sampling, race, age category (> and < median), sex and sequencing batch. Only groups with and without the primary tumor present at time of sampling show a statistically significant difference (p-value = 0.046) in area under the curve (AUC) based on the DeLong test performed on the median model. CRC: colorectal adenocarcinoma; AA: appendiceal adenocarcinoma; CI: confidence interval; SD: standard deviation.

**Supplementary Figure S6:**
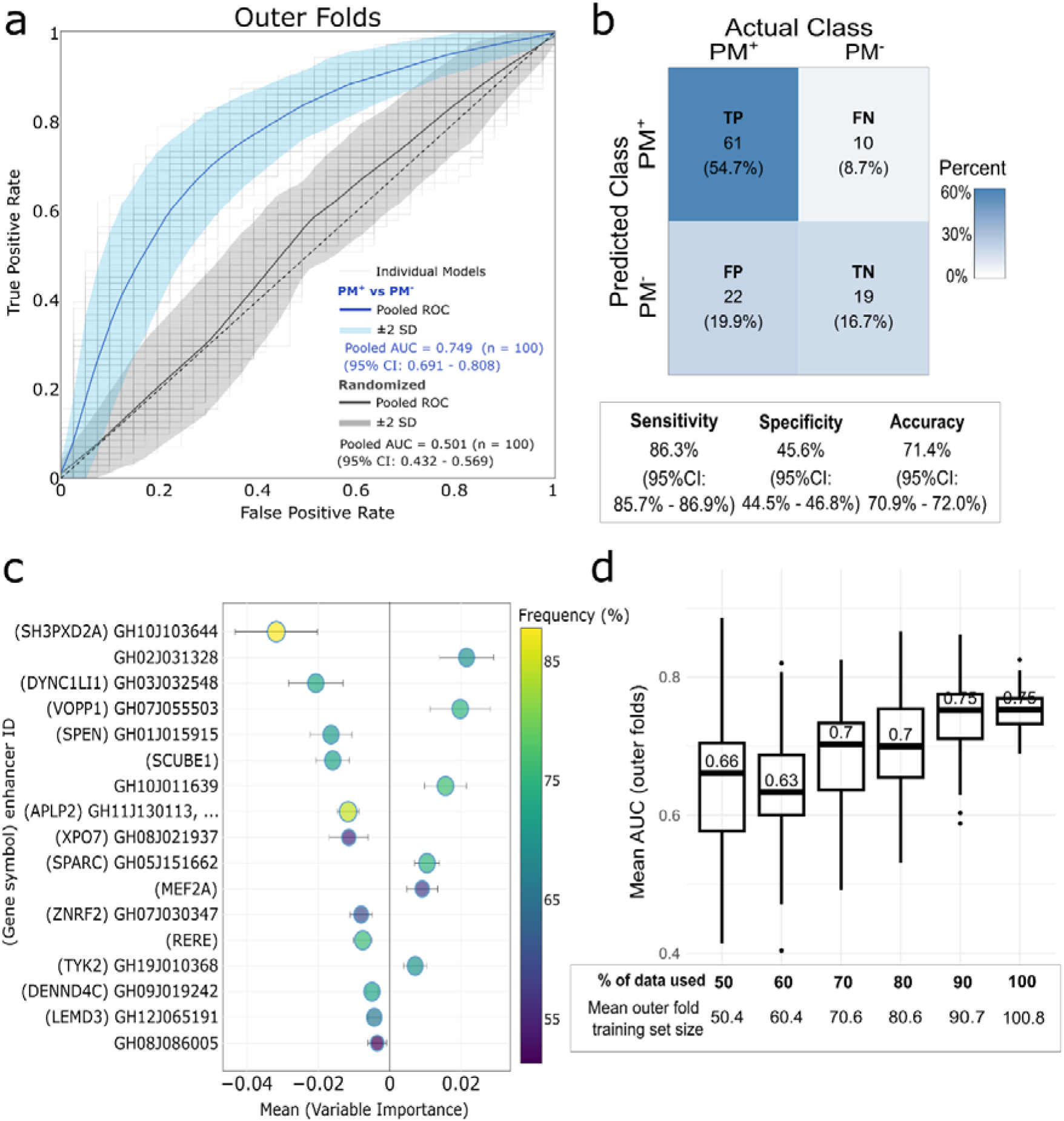
Performance of sparse models predictive of PM status. **a** Receiver operating characteristic (ROC) curves for outer fold test sets from 10-fold nested cross-validation (CV) of elastic net regularization models trained on top 50 features retained from a random forest (RF) filter (sparse model configuration), predictive of peritoneal metastasis (PM) status. Pooled area under the curve (AUC) of 0.749 (95% CI 0.691-0.808) (n = 1,000) was reached relative to 0.501 (95% CI 0.432-0.569) (n = 1,000) for negative control with randomized PM status labels indicating results are likely not due to chance nor overfitting. **b** Pooled confusion matrix for all outer-fold test sets, for each run with a different random seed (n = 100), for models predictive of PM status with sparse model configuration, showing higher sensitivity of 86.3% relative to specificity of 45.6%, and an accuracy of 71.4% (95% CI 70.9-72.0%). **c** Variable importance plot showing mean DhMR coefficients of features occurring in > 50% of models (n = 1,000) predictive of PM status. Features show highly stable model coefficient scores, as shown by error bars representing pooled standard error of the mean (SEM). A mean coefficient > 0 indicates 5hmC enrichment in these genomic regions is associated with an increased probability of cancer or peritoneal metastasis. **d** Box plots displaying mean outer fold AUC relative to the number of samples used for model training and evaluation showing an increase in model accuracy with increasing amounts of training data (20 random seeds evaluated per configuration, resulting in n=200 outer-fold models for each configuration).

**Supplementary Figure S7:**
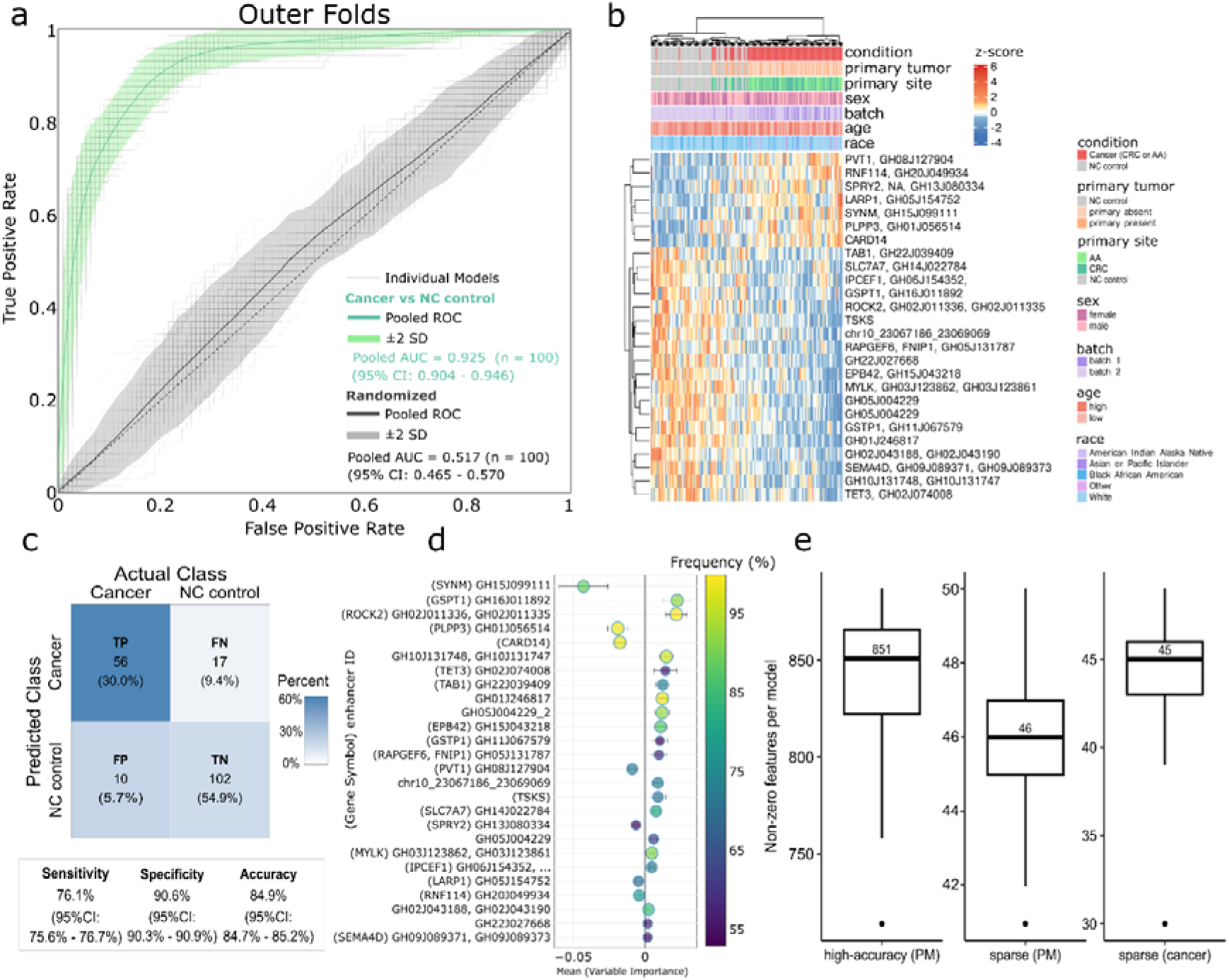
Detailed performance of model predictive of cancer status relative to non-cancer controls. **a** Receiver operating characteristic (ROC) curves for outer folds from 10-fold cross-validation (CV) of elastic net regularization models trained on top 50 features (sparse configuration) retained from random forest (RF) filter, predictive of CRC or AA cancer status relative to non-cancer (NC) controls. Average area under the curve (AUC) of 0.925 (95% CI 0.904-0.946) (n = 1,000) was reached relative to 0.517 (95% CI 0.465-0.570) (n = 1,000) for negative control with randomized PM status labels. **b** Heatmap showing sample clustering by cancer status based on the z-score of 26 hydroxymethylated regions (hMRs) from (**d**). Samples and hMRs were clustered using the Ward.D2 Euclidean distance metric. **c** Pooled confusion matrix for all outer-fold test sets, for each run with a different random seed (n = 100), for models predictive of CRC or AA cancer status reaching mean sensitivity of 76.1% relative to specificity of 90.6%, and an accuracy of 84.9% (95% CI 84.7-85.2%). **d** Variable importance plot showing mean DhMR coefficients of features occurring in > 50% of models (n = 1,000 outer-fold models) predictive of CRC or AA cancer status. Features show highly stable model coefficient scores, as shown by error bars representing pooled standard error of the mean (SEM). A mean coefficient > 0 indicates 5hmC enrichment in these genomic regions is associated with an increased probability of cancer status. **e** Box plots displaying mean number of hMR features with non-zero coefficients per model, for sparse and high-accuracy parameters for models predictive of PM status (retaining 50 and 5,000 top features after RF respectively, as shown in Figure 3c), and for sparse model parameters for models predictive of CRC or AA cancer status. CRC: colorectal adenocarcinoma; AA: appendiceal adenocarcinoma.

**Supplementary Figure S8:**
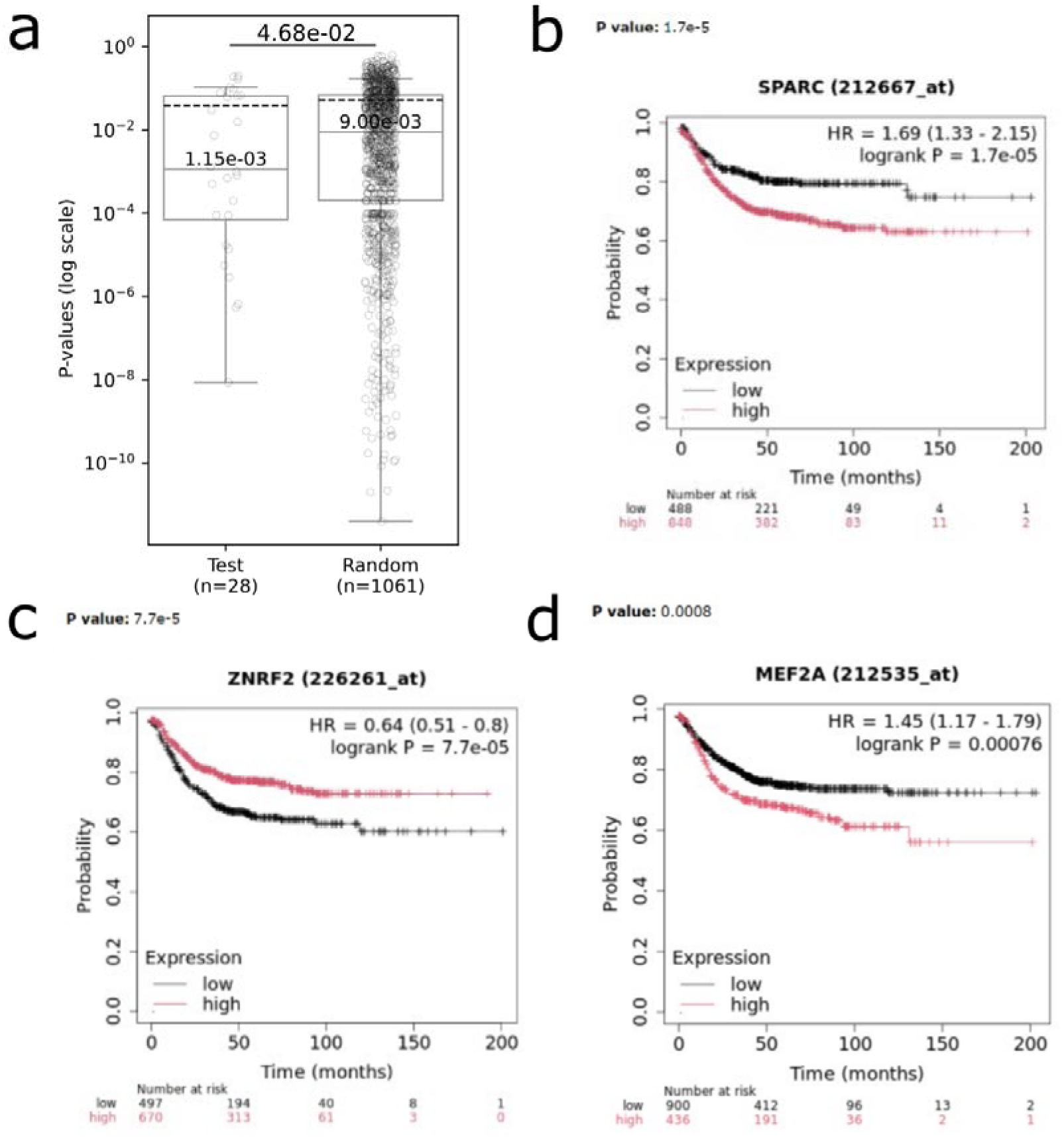
Probability of differences in RFS for genes with high occurrence (>99%) PM predictive models. **a** Genes associated with hMRs occurring in >99% of high-accuracy models predictive of PM status were found to have statistically significant (p-value = 0.041) association with relapse-free survival (RFS) differences in TCGA CRC patients (n = 1,167). “Test” group represents the set of genes associated with PM-predictive hMRs (n = 28), “Random” represents the set of genes drawn at random (n = 1,061) from all human gene symbols. The test set had a median p-value of RFS difference of 1.15e-3 relative to 9.00e-3 for the random set. Difference in medians was evaluated with a permutation test with 10,000 iterations. **b–c** Two select genes associated with regulatory regions based on genomic position (within gene body) or enhancer regulatory targets (MEF2A, GMPS, SPARC and ZNRF2) show statistically significant differences in 5-year survival when stratified based on high / low mRNA expression from CRC patients in TCGA database (n = 1,167).

